# TCF12 controls oligodendroglial cell proliferation and regulates signaling pathways conserved in gliomas

**DOI:** 10.1101/2021.07.26.453859

**Authors:** Sofia Archontidi, Corentine Marie, Beata Gyorgy, Justine Guegan, Marc Sanson, Carlos Parras, Emmanuelle Huillard

## Abstract

Diffuse gliomas are primary brain tumors originating from the transformation of glial cells. In particular, oligodendrocyte precursor cells constitute the major tumor-amplifying population in the gliomagenic process. We previously identified the *TCF12* gene, encoding a transcription factor of the E protein family, as being recurrently mutated in oligodendrogliomas. In this study, we sought to understand the function of TCF12 in oligodendroglial cells, the glioma lineage of origin. We first describe TCF12 mRNA and protein expression pattern in oligodendroglial development in the mouse brain. Second, by TCF12 genome wide chromatin profiling in oligodendroglial cells, we show that TCF12 binds active promoters of genes involved in proliferation, translation/ribosomes, and pathways involved in oligodendrocyte development and cancer. Finally, we perform OPC-specific *Tcf12* inactivation *in vivo* and demonstrate by immunofluorescence and transcriptomic analyses that TCF12 is transiently required for OPC proliferation but dispensable for oligodendrocyte differentiation. We further show that *Tcf12* inactivation results in deregulation of biological processes that are also altered in oligodendrogliomas. Together, our data suggest that TCF12 directly regulates transcriptional programs in oligodendroglia development that are relevant in a glioma context.

## Introduction

Diffuse gliomas are the most prevalent malignant primary brain tumors in adults (Ostrom et al., 2017). Gliomas are classified according to their histological and genomic features, including the presence of mutations in isocitrate dehydrogenase genes (mostly in *IDH1*) and the status of the loss of the chromosomal arms 1p and 19q (termed as *1p/19q* co-deletion) (Louis et al., 2016). The three main glioma entities are oligodendrogliomas (*IDH* mutated, *1p/19q* co-deleted), astrocytomas (*IDH* mutated, *1p/19q* intact) and glioblastomas (also termed GBM, *IDH* wild type, *1p/19q* intact), with the latter being most aggressive.

Gliomas contain cells with features of glial cells and neural stem/progenitor cells (NSC/NPCs) that are endowed with proliferative capacity (Bielle et al., 2017; Neftel et al., 2019; Tirosh et al., 2016). Furthermore, mounting evidence from genetically engineered mouse models indicate that cells with characteristics of NSC/NPCs, or oligodendrocyte precursor cells (OPCs) are potential cells of origin and responsible for tumor amplification in gliomas (Alcantara Llaguno et al., 2009; C. Liu et al., 2011; Persson et al., 2010; Weng et al., 2019). Thus, deciphering how alterations found in gliomas impact oligodendroglial lineage cells is critical to understand the cellular and molecular mechanisms underlying tumor development and progression.

In the forebrain, OPCs are generated from NSC/NPCs at both embryonic and postnatal stages, and after proliferating and populating the brain, they start to differentiate into myelin-producing oligodendrocytes (OLs) around the second week after birth (Kessaris et al., 2006; Shen et al., 2021). Recent studies have described oligodendroglial differentiation as a continuum, starting from OPCs, followed sequentially by committed oligodendrocyte progenitors (COPs), newly formed OLs (NFOLs), myelin forming OLs (MFOLs) and finally by several populations of mature, axon ensheathing, OLs (MOLs) (Marques et al., 2016, 2018). OPCs constitute the major proliferating cell type the adult central nervous system (CNS), acting as source for differentiated OLs while maintaining the pool of cells available for differentiation (Dawson et al., 2003). OPC proliferation and differentiation properties are subjected to a strict, finely tuned regulation by a complex network of signaling, transcription and epigenetic factors (reviewed in (Parras et al., 2020; Sock & Wegner, 2019).

Transcription factor 12 (*TCF12*, also called *HTF4* or *HEB*) is a member the E protein family, a subclass of the basic helix-loop-helix (bHLH) protein family that includes TCF3 (E2A) and TCF4 (E2-2) (Massari & Murre, 2000). Functionally, TCF12 has been reported to regulate the differentiation of lymphocytes (Emmanuel et al., 2018; Jones-Mason et al., 2012; Wojciechowski et al., 2007), the formation of germ layers from embryonic stem cells (Li et al., 2017; Yi et al., 2020; Yoon et al., 2015), osteoblast differentiation (Yi et al., 2017) and cranial suture development (Sharma et al., 2013). In the CNS, TCF12 is implicated in the development of midbrain dopaminergic neurons (Mesman & Smidt, 2017) but its roles in the oligodendroglial lineage are not known. Early studies had demonstrated that *TCF12* is expressed in oligodendrogliomas and astrocytomas (Riemenschneider et al., 2004). Furthermore, we and others previously reported that *TCF12* is mutated in oligodendrogliomas (Labreche et al., 2015; Aihara et al., 2017; Suzuki et al., 2015). In particular we found that *TCF12* mutations resulted in reduced transcriptional activity and were associated with aggressive tumor features (Labreche et al., 2015). However, how *TCF12* mutations functionally impact the development of glioma cells of origin are not understood.

In this study, we explored the functions of TCF12 in oligodendroglia and gliomas, using chromatin binding profiling, transcriptomic and genomic analyses as well as genetic mouse models of *Tcf12* conditional deletion. We report that TCF12 positively regulates OPC proliferation *in vivo* and further suggest putative implications that can be conserved in a glioma context.

## Methods

### Mice

Mice were housed, bred and treated in an authorized facility (agreement number A751319). All protocols and procedures involving mice were ethically reviewed and approved by the local ethics committee and the French Ministry of Research and Higher Education (approval number APAFIS#20939-2020052811427837). Swiss (RjOrl:SWISS) mice were obtained from Janvier Laboratories (France). *Tcf12flox* mice (*Tcf12*^*tm3Zhu*^ *Tcf3*^*tm4Zhu*^/J; (Wojciechowski et al., 2007)) were obtained from Jackson Laboratory and maintained as heterozygotes (*Tcf12*^*flox/wt*^) by crossing them with C57BL/6JRj mice (Janvier Laboratories). These mice were bred to *PDGFRα::CreER*^*T*^ (Kang et al., 2010) and *Rosa26*^*LSL-YFP*^ (Srinivas et al., 2001) to generate *PDGFRα::CreER*^*T/wt*^; *Rosa26*^*LSL-YFP/ LSL-YFP*^; *Tcf12*^*flox/wt*^ males that were subsequently bred with *Tcf12*^*flox/wt*^ females. Both male and female animals were used for the experiments. Genotyping was performed using standard protocols (sequences of genotyping primers are available upon request). To induce Cre-mediated DNA recombination, *PDGFRα::CreER*^*T*^*;Rosa26*^*LSL-YFP*^*;Tcf12*^*flox*^ mice were injected subcutaneously with 100μg/g of tamoxifen (T5648, Sigma; dissolved in corn oil; stock concentration of 20mg/mL) at P13, once a day for three consecutive days.

### Immunofluorescence on frozen sections

Mice were sedated by xylazine (Rompun, 10-15mg/kg) and euthanized by intraperitoneal injection of sodium pentobarbital (Euthasol, 140mg /Kg). Mice were intracardially perfused with NaCl 0.9% followed by PFA 2% (Electron Microscopy Sciences 15713, diluted in PBS). Brains were post-fixed in PFA 2% and cryoprotected in 20% sucrose overnight at 4°C. The following day, brains were embedded in OCT (16-004004, Tissue-Tek), frozen in dry-ice-chilled isopentane and stored at -80 °C. Fourteen-micron cryosections were obtained using a cryostat (Leica). Cryosections were air-dried at room temperature (RT) and then incubated with blocking solution, containing 10% normal goat serum (NGS) in 0.3 % Triton X-100 (Sigma) in PBS (PBS-Triton 0.3%), for 1 hour at RT. Subsequently, sections were incubated with the primary antibodies either at 4°C overnight or at RT for 2 hours. After three washes with PBS, sections were incubated with fluorophore-labeled secondary antibodies for 1 hour at RT. Both primary and secondary antibodies were diluted in blocking solution (10% NGS in 0.3 % PBS-Triton). After three washes with PBS, sections were incubated in DAPI solution (300nM, Invitrogen D3571) for nuclear counterstaining, for 10 minutes at RT, and mounted using Fluoromount™ Aqueous Mounting Medium (Sigma-Aldrich). Slides were kept at 4°C until image acquisition. Immunofluorescent staining of fixed cells was performed as described above, except that a solution of 0.05 % Tween 20 (P2287 Merck) in PBS was used instead of 0.3 % Triton -PBS. The references of primary and secondary antibodies are given in Supplementary Information.

### Image acquisition and quantification

Images were captured using either a Leica Sp8x Confocal Microscope or a Zeiss wide-field fluorescent microscope (equipped with an Apotome system). Image processing and analysis was performed on ZEN 2.0 blue edition software (Zeiss) or Fiji (Schindelin et al., 2012). Images were generated as maximum intensity projections (MIPs) of the entire imaging depth. Cell counting was done with Fiji’s Cell Counter plugin on MIPs.

### Magnetic Activated Cell Sorting (MACSorting) of oligodendroglial cells

Mice (P12-P16) were euthanized with CO_2_ followed by immediate decapitation. Brains were rinsed in cold PBS and the dorsal region containing the cortex and corpus callosum was harvested. Dissociation was performed using the gentleMACS Octo Dissociator with Heaters (program 37C _NTDK _1, Miltenyi Biotec) with the appropriate tubes (gentleMACS C Tubes, 130-093-237, Miltenyi Biotec) and an enzymatic mix composed of 0.46 mg/mL papain (WOLS03126, Worthington), 0.1 mg/mL DNAse (WOLS02139, Worthington) and 0.124 mg/mL L-cysteine (C7880, Sigma) and HBSS 1X supplied with Ca^2+^ and Mg^2+^. Upon dissociation, the suspension was filtered (130-110-916, Miltenyi Biotec), the cells were further resuspended in 5 volumes of cold HBSS and centrifuged at 300g for 10 minutes at 4°C. Additionally, a debris removal step was performed using a Debris Removal Solution (130-109-398, Miltenyi Biotec). Subsequently, cells were incubated with anti-O4 Microbeads (130-094-543, Miltenyi Biotec) and magnetic separation was done using Multi-24 Column Blocks and the MultiMACS Cell24 Separator Plus (130-095-692 and130-098-637, Miltenyi Biotec). O4^+^ cells were collected in BSA 0.5 % in PBS solution and counted. To characterize the sorted cell population, part of cell suspension was plated poly-ornithine coated (P4957, Sigma) and cultured for 2 hours before 4% PFA fixation as described in (Marie et al., 2018).

### Chromatin immunoprecipitation followed by sequencing (ChIP-Seq)

We used NPCs (cultures established from wild type Swiss neonatal mice) or acutely isolated O4^+^ MACSorted cells (harvested from P11-P12 wild type Swiss mice). We used 4.10^6^ cells for immunoprecipitation with TCF12 antibody and 10^6^ cells for immunoprecipitation with each histone (H3K27Ac, H3K4me3, H3K27me3) antibody. Cells were fixed in PFA 1% for 10 minutes at RT. Fixation was quenched with 125 mM glycine (G8898, Sigma) for 5 minutes and cells were washed in cold PBS supplied with protease inhibitor cocktail (11873580001, Roche). Cells were stored as dry cell pellet at -80°C until further processed. The next steps were performed using iDeal ChIP-Seq kit for Transcription Factors (C01010055, Diagenode). Briefly, cells were lysed, and chromatin was sheared using a Bioruptor Pico sonicator (10 sonication cycles 30’’ ON/ 30’’ OFF, Diagenode). Sheared chromatin was incubated under constant rotation at 4°C O/N with Protein A-coated magnetic beads, coupled with rabbit anti-TCF12 antibody (5 μg, SAB3500566, Sigma). Elution, cross-link reversal and DNA purification steps were performed according to the manufacturer’s protocol (Diagenode). Input (non-immunoprecipitated sheared chromatin) was used as control. Protocols for H3K27Ac, H3K4me3, H3K27me3, are described elsewhere (C Marie and C Parras, unpublished data). The ChIP-Seq libraries were prepared using TruSeq ChIP library preparation kit (ILLUMINA) and sequenced with a Nextseq 500 platform (ILLUMINA, 57 10^6^ of 75 bp pair-end reads per sample). Sequenced datasets were processed with the Galaxy suite (https://usegalaxy.org/). Reads were trimmed using Cutadapt and Trimmomatic. Data was aligned to the mouse mm10 genome, using Bowtie2. PCR-derived duplicates were removed using PICARD MarkDuplicates and blacklisted regions were removed with blacklist. Bigwig coverage files were generated with bamCoverage and peak calling was performed using MACS (Model-based Analysis of ChIP-Seq) with options: --keep-dup 1, --narrow, --nomodel and filtered according to the following criteria: (i) length>=100bp and (ii) p-value<=5%. The Input for each individual experiment was used as control. Representation of the data was done using IGV browser (https://software.broadinstitute.org/software/igv/, (Robinson et al., 2011)). Overlapping, region annotation and correlations were done using Genomatix (www.genomatix.de). Gene set enrichment analyses were done using Enrichr (https://maayanlab.cloud/Enrichr/,(Chen et al., 2013)). “Promoters” correspond to regions 1000bp upstream of transcription start site (TSS) and 10bp downstream of TSS (Genomatix). “Enhancers” correspond to the regions associated with the presence of histone marks outside promoters.

### RNA extraction, RT-qPCR and sequencing

mRNAs of O4^+^ MACSorted cells were extracted using the Macherey-Nagel NucleoSpin RNA XS kit (740902.50, Macherey-Nagel) and quantified with Nanodrop spectrophotometer. RNAs were reverse transcribed to cDNA using the Maxima 1str cDNA Synth Kit (K1642, LifeTechnologies). Quantitative PCR was performed using LightCycler 480 SYBR Green I Master Mix (4707516001, Roche) on a LightCycler® 96 thermocycler. Samples were run in replicates (duplicates or triplicates). Primers details are listed in the Supplementary Information. *Gapdh* and *Tbp* genes were used for normalization. Analyses were performed using the delta-delta Cq method.

For RNA sequencing, RNA-Seq libraries were prepared using the NEBNext Ultra II Directional RNA Library Prep Kit (NEB) and sequenced with the Novaseq 6000 platform (ILLUMINA, 32*10^6^ 100bp pair-end reads per sample). Quality of raw data was evaluated with FastQC. Poor quality sequences were trimmed or removed with fastp tool, with default parameters, to retain only good quality paired reads. Illumina DRAGEN bio-IT Plateform (v3.6.3) was used for mapping on mm10 reference genome and quantification with gencode vM25 annotation gtf file. Library orientation, library composition and coverage along transcripts were checked with Picard tools. Subsequent analyses were conducted with R software. Data were normalized with edgeR (v3.28.0) bioconductor packages, prior to differential analysis with glm framework likelihood ratio test from edgeR package workflow. Multiple hypothesis adjusted p-values were calculated with the Benjamini-Hochberg procedure to control False Discovery Rate (FDR). Finally, enrichment analysis was conducted with clusterProfiler R package (v3.14.3) using Gene Set Enrichment Analysis (GSEA), on hand curated collections and on collections of the MSigDB. For the differential expression analyses, low expressed genes were filtered, sex was used as covariable and the cut-offs applied were: FDR < 0.05 and log2FC > 0.5.

### Statistical analysis

Data were plotted and analyzed using MS Excel, GraphPad Prism 8 or R Studio, unless otherwise specified. In bar graphs, data are presented as mean + standard error of the mean (SEM). Points indicate independent biological samples (n) of the same genotype. Statistical tests used are specified in the figure legends.

## Results

### *TCF12* is altered in gliomas

To extend previous studies and obtain a broad view of the type and distribution of *TCF12* alterations in gliomas, we queried the “Lower Grade Glioma” (LGG, grades II-III) and “Glioblastoma Multiforme” (GBM, grade IV) data sets of the TCGA PanCancer Atlas (containing data for 514 and 592 tumors respectively). By interrogating those datasets for mutations and copy number alterations, we found that *TCF12* alterations are present in all glioma types (Supplementary Figure 1A). *TCF12* is altered in approximately 27% and 18% of the GBM and LGG respectively (Figure 1A). In both tumor types, 70-80% of *TCF12* alterations correspond to heterozygous losses (Figure 1B). We next compared the occurrence of *TCF12* alterations with other common glioma alterations. We found that *TCF12* alterations co-occur with the most common glioma alterations (*TP53, PTEN, NF1, EGFR, CDKN2A*) but do not co-occur significantly with *IDH1* or *CIC* alterations (Supplementary Figure 1A and 1B; Supplementary Table 1), further indicating that TCF12 alterations are not specific to a glioma type. We next analyzed whether *TCF12* alterations were associated with patient clinical outcome. We did not detect an association between *TCF12* alterations and patient overall survival (Figure 1C and 1D). However, when considering *TCF12* high and low expression levels (Supplementary Table 2), we noted that patients with higher *TCF12* expression had a better survival compared to *TCF12*-low patients, both in GBM and LGG (Figure 1E and 1F), in line with a recently published study (Noorani et al., 2020). Together, these data suggest a broad and tumor suppressive function for TCF12 in gliomas.

**Figure 1:**
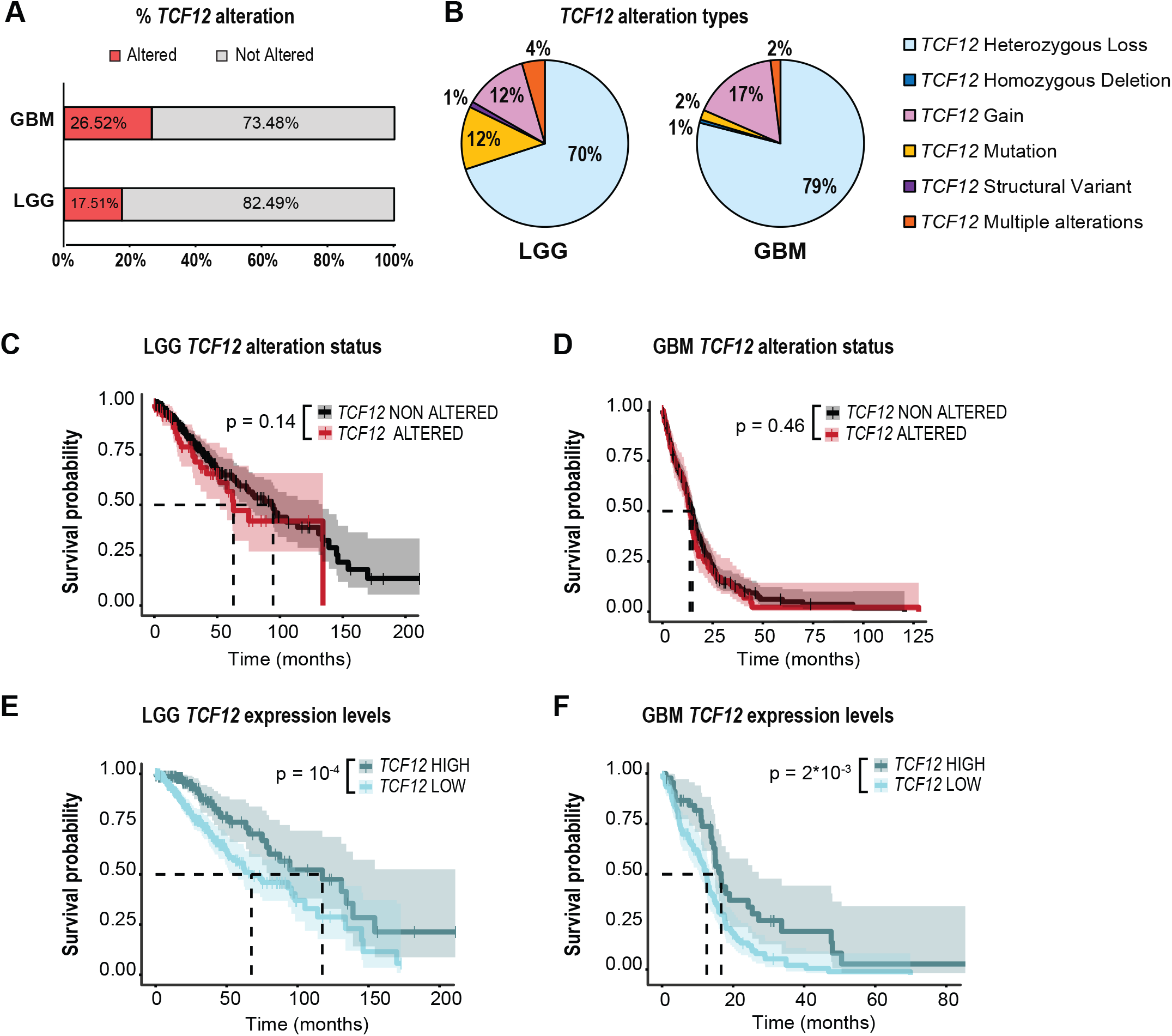
*TCF12* is altered in gliomas. **A**. Bar graph showing the percentage of *TCF12* altered samples in the glioblastoma (GBM, 592 samples) and lower grade glioma (LGG, 514 samples) cohorts from The Cancer Genome Atlas (TCGA). **B**. Pie charts illustrating the types of alterations in *TCF12* altered gliomas of Figure 1A. **C-D**. Kaplan-Meier survival curves of TCGA LGGs (n=77 altered, n=418 non altered, C) and GBMs (n=95 altered, n=263 non altered, D) comparing *TCF12* altered vs non altered groups. **E-F**. Kaplan-Meier survival curves of TCGA LGGs (n=154 high, n=359 low, E) and GBMs (n=46 high, n=106 low, E) comparing high or low *TCF12* mRNA expression.

### TCF12 is expressed in oligodendroglial cells

Given that cells with features of OPCs constitute a major tumor driving population in gliomas (C. Liu et al., 2011; Weng et al., 2019), we thus explored TCF12 function in these cells. In order to characterize *Tcf12* expression across the oligodendrocyte lineage, we first interrogated a bulk RNA-Seq transcriptome database of mouse and human cerebral cortex cell types (Zhang *et al* 2014), finding that *Tcf12* is expressed in neurons, astrocytes and in oligodendroglial cells in the mouse and human brain, with higher expression levels in oligodendroglia (Figure 2A). We next processed and integrated single cell transcriptomic datasets of embryonic and postnatal mouse oligodendroglial lineage cells (Marques et al., 2016; 2018) to further explore *Tcf12* expression across the oligodendrocyte (OL) lineage (Supplementary Figure 2A and 2B). Analysis of this data revealed *Tcf12* expression in NSCs, NPCs, and OL lineage cells, with *Tcf12* expression being higher in early progenitor cells and differentiating OLs, compared to mature OLs (Figure 2B,C). Given the functional compensation among E-proteins suggested in neuroglial and other lineages (Ravanpay & Olson, 2008; Wedel et al., 2020; Zhuang et al., 1998), we also analyzed the expression of *Tcf3* and *Tcf4* in OL lineage cells. While*Tcf3* was expressed in few cells and at quite low levels, *Tcf4* was expressed in a higher percentage of NSCs, NPCs, and oligodendroglia than *Tcf12*, with a peak of expression in NPCs and OPCs (Supplementary 2C). Finally, we validated the presence of TCF12 protein in oligodendroglial cells in wild type mice by performing immunofluorescence of TCF12 along with PDGFRa (identifying OPCs), and the combination of CC1 and OLIG1 (identifying OLs, Figure 2D). We show that TCF12 protein is present in OPCs and differentiating OLs in the mouse corpus callosum, at postnatal and adult stages (Figure 2E-F).

**Figure 2:**
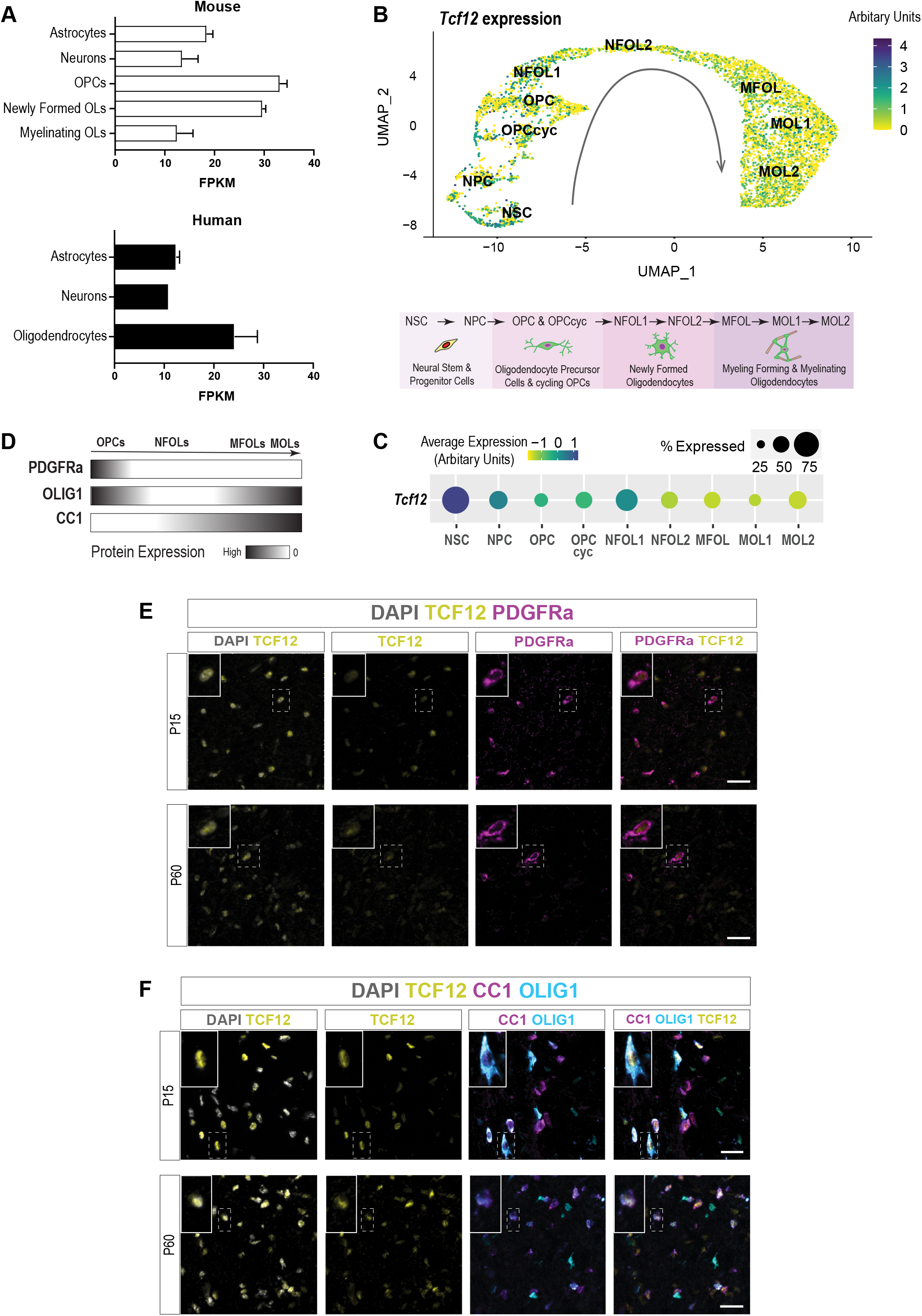
TCF12 is expressed in oligodendroglial cells. **A**. Bar graphs showing the expression of *TCF12* mRNA in astrocytes, neurons and oligodendroglial cells in mouse (top) and human (bottom) brain (data extracted from https://www.brainrnaseq.org/) **B**. Reconstructed UMAP representation illustrating *Tcf12* expression across oligodendroglia differentiation (direction of differentiation indicated by the arrow, yellow = low expression and dark blue = high expression, related to Supplementary Figure 2 A-B). **C**. Dot plot showing mean *Tcf12* expression and the percentage of *Tcf12*-expressing cells in each cluster shown in Supplementary Figure 2B. **D**. Schematic representation of the expression of selected OPC and OL markers (according to (Nakatani et al., 2013)). **E-F**. Immunostaining of TCF12 with PDGFRa (OPCs, E) or CC1 and OLIG1 (OLs, F) in the corpus callosum of postnatal (P15) and adult (P60) wild type mice. Scale bars, 20μm. Insets represent 60% magnifications of the cells highlighted dash-lined squares. OPCs = oligodendrocyte precursor cells, OLs = oligodendrocytes.

### TCF12 primarily occupies active promoter regions in oligodendroglial cells

To explore the potential roles mediated by TCF12 in the oligodendrocyte lineage, we next sought to identify putative TCF12 gene targets and pathways in oligodendroglial cells, by generating the chromatin binding profiles of TCF12 in NSC/NPCs and oligodendroglia. To do so, we first performed magnetic-activated cell sorting (MACS) using O4 antibodies (recognizing OPCs/OLs, (Dincman et al., 2012) to purify oligodendroglial cells from wild type postnatal day 12 (P12) mice, obtaining a population composed of ∼30% OPCs and ∼60% OLs (thereafter termed “OPCs/OLs”, Supplementary Figure 3A-B). To assess cell type-specific and overlapping targets for TCF12, we also prepared neurosphere cultures of neural progenitor cells (thereafter termed “NPCs”, Supplementary Figure 3A). We then performed chromatin immunoprecipitation followed by DNA sequencing (ChIP-Seq) using antibodies directed against TCF12 and histone marks defining status of regulatory elements as active, poised and repressed. Elements harboring H3K27Ac/H3K4me3 were considered as active, H3K4me3 alone as poised, and H3K27me3 as repressed (Rada-Iglesias et al., 2011). Peak calling identified 18405 TCF12 binding sites in NPCs and 28654 in OPCs/OLs that were associated with 6947 and 15670 genes, respectively (Figure 3A, Supplementary Table 3). Only 3032 genes were shared between OPCs/OLs and NPCs, suggesting that TCF12 binding is cell type specific (Figure 3A). Analysis of the distribution of TCF12 binding sites over genomic regions (promoters, exons, introns, and intragenic regions) indicated that TCF12 binding in OPCs/OLs was particularly enriched in promoters, representing 42% of bound regions, compared to only 7% in NPCs (Figure 3B). Visualizing the genomic distribution of TCF12 binding sites further revealed a strong enrichment of TCF12 binding in the proximity of promoters in OPCs/OLs but not in NPCs, whereas TCF12 bound the vicinity of enhancers, defined as regions harboring a histone mark located away from a transcription start site, in both NPCs and OPCs/OLs (Figure 3C). We then characterized whether the regulatory regions bound by TCF12 were active, poised or repressed and found that promoter regions bound by TCF12 were mainly active both in OPCs/OLs and NPCs (52% active, 12% poised, and 6% repressed in OPCs/OLs, and 63% active, 23% poised, and 5% repressed in NPCs), while enhancer regions bound by TCF12 binding corresponded mostly to poised enhancers (52% in OPCs/OLs and 69% in NPCs) (Figure 3D and Supplementary Figure 3C, Supplementary Table 3). Altogether, our findings of TCF12 occupancy in gene regulatory regions enriched in active or poised chromatin marks in both OPCs/OLs and NPCs, suggest that it directly activates gene expression in both cell types. Notably, the promoters of genes known as markers of OPCs (*Pdgfra*) and OLs (*Itpr2, Mbp*), as well as proliferation markers (*Mki67, Cdkn1a*), were bound by TCF12 in OPCs/OLs and displayed active histone marks (Figure 3E). Finally, we performed enrichment analysis of the genes associated with TCF12 binding at active promoters in OPCs/OLs. This analysis revealed enrichment of pathways related to proliferation, translation/ribosomes, proteasome, and signaling pathways involved in oligodendrocyte development and cancer (such as MYC, TGFß, PI3K/AKT/mTOR, WNT-beta catenin, p53, Notch) (Figure 3F; Supplementary Figure 3D; Supplementary Table 4). Therefore, all together, these data suggest that TCF12 regulates oligodendrocyte development, by positively regulating gene expression.

**Figure 3:**
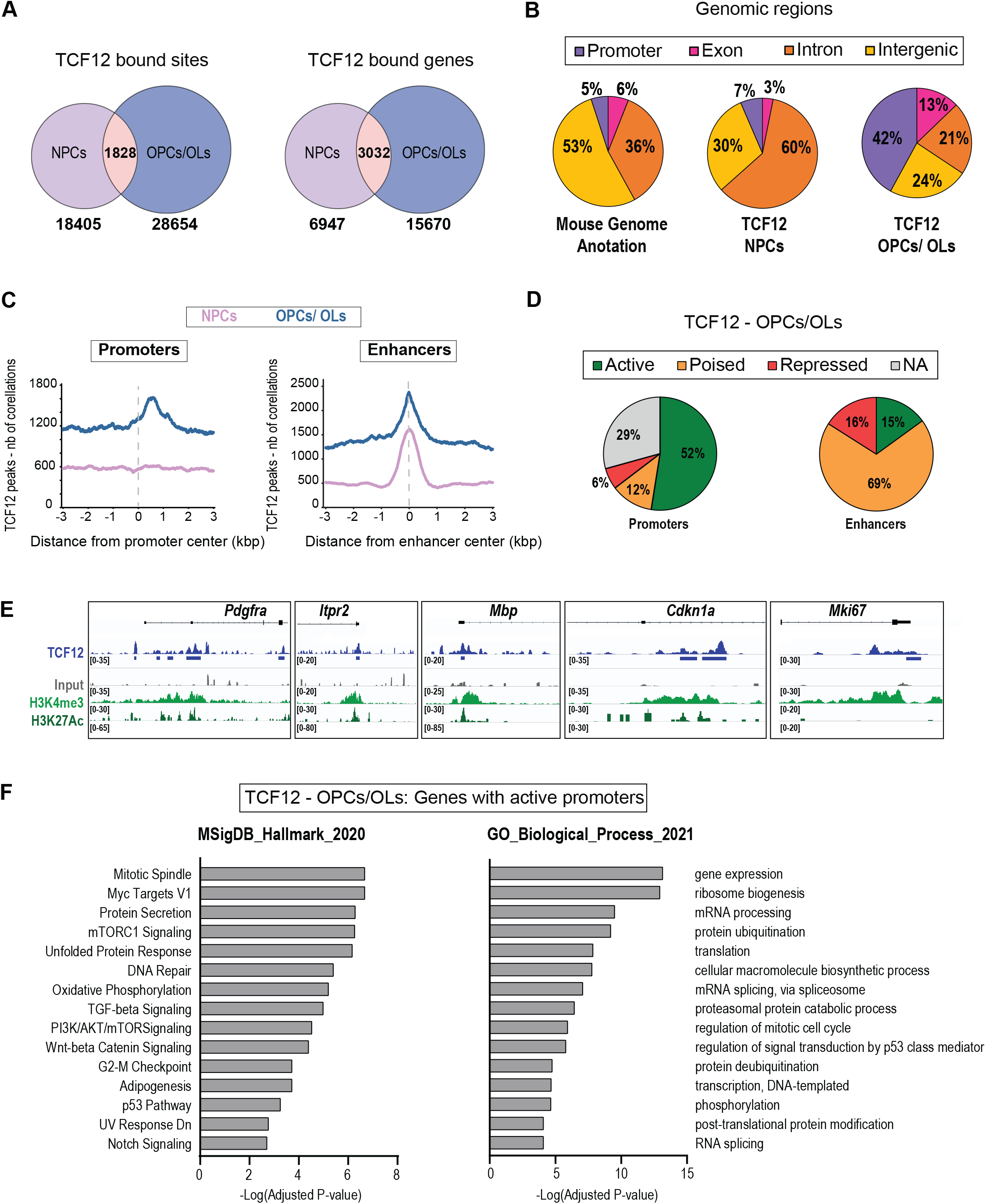
TCF12 mainly occupies active promoter regions in oligodendroglial cells. **A**. Venn diagrams illustrating the overlap of TCF12 bound sites and genes in NPCs and OPCs/OLs. **B**. Graphs depicting the annotation of TCF12 bound sites in NPCs (middle) and OPCs/OLs (right) with genomic regions compared to the region representation in the genome (left). **C**. Graphs showing the number of correlations of TCF12 peaks in NPCs (purple line) and OPCs/OLs (blue line) compared the central position of promoter (left) and enhancer (right) regions. **D**. Pie charts showing the distribution of TCF12 bound sites in promoter (left) and enhancer (right) regions in OPCs/OLs in association with epigenetic marks: H3K4me3 and H3K27Ac = active, H3K4me = poised, H3K27me3= repressed, NA=no epigenetic mark. **E**. Representative ChIP-Seq tracks for TCF12, input control and active epigenetic marks (H3K4me3 and H3K27Ac) in *Pdgfra, Itpr2, Mbp, Ki67* and *Cdkn1a* in promoter regions in OPCs/OLs. **F**. Bar plots showing significantly enriched terms and pathways in the TCF12 bound genes with active promoters in OPCs/OLs (6596 genes). For the “MSigDB_Hallmark_2020” library, the top 15 most significant gene sets are displayed. For the “GO_Biological_Process_2021” library, 15 among the top 50 most significant gene sets are displayed.

### TCF12 inactivation in OPCs *in vivo* leads to proliferation defects

To further characterize the roles of TCF12 in oligodendrocyte lineage, we generated an inducible *Tcf12* knockout mouse model. To do so, we combined a *Tcf12*^*flox*^ mice having loxP sites flanking the exons encoding the bHLH domain (Mesman & Smidt, 2017; Wojciechowski et al., 2007), with mice carrying *Pdgfra::CreER*^*T*^ driver (Kang et al., 2010) and *YFP* inducible reporter (Srinivas et al., 2001). Thus, in this *Pdgfra::CreER*^*T*^*;Rosa26*^*LSL-YFP*^*;Tcf12*^*flox*^ model, TCF12 inactivation can be induced specifically in OPCs upon tamoxifen-dependent Cre-mediated recombination and the cell fates of *Tcf12*-mutant cells can be traced by the YFP reporter. In this model, we compared mutant heterozygous (*Tcf12*^*het*^) and homozygous (*Tcf12*^*hom*^) animals with intact Tcf12 (*Tcf12*^*ctrl*^) littermates (Supplementary Figure 4A-B). As a first step, we administrated tamoxifen at P13, at the peak of oligodendrocyte differentiation in the corpus callosum, and performed our analysis at P16 to focus on the immediate effects following *Tcf12* inactivation (Figure 4A). We first validated the efficacy of Cre-mediated recombination of the *Tcf12*^*flox*^ allele by quantifying mutated *Tcf12* transcripts by RT-qPCR on OPCs/OLs purified by O4^+^ magnetic sorting from P16 cortices. We observed a 25% to 50% decrease in *Tcf12* transcript levels in *Tcf12*^*het*^ and *Tcf12*^*hom*^ animals respectively, compared to *Tcf12*^*ctrl*^ (Supplementary Figure 4D), which is consistent with ∼60% of O4+ cells being recombined (YFP^+^) at the time of analysis (Supplementary Figure 4E). We next asked whether *Tcf12* inactivation affected OPC proliferation and differentiation properties by performing combined immunostaining with antibodies against GFP to detect the recombined (YFP^+^) cells, PDGFRα/Ki67 to label OPCs and their proliferative status, and CC1/OLIG1 to label different OL stages (Figure 4B, C). Remarkably, while the density of recombined OPCs (PDGFRα^+^GFP^+^/mm^2^) (Figure 4D) and the fraction of recombined OPCs (PDGFRα^+^GFP^+^/PDGFRα^+^ and PDGFRα^+^GFP^+^/ GFP^+^) (Figure 4E,F) remained unchanged in the corpus callosum across the different genotypes, the fraction of proliferating OPCs was reduced by two-fold in *Tcf12*^*hom*^ animals (28.2% ± 3.94% in *Tcf12* ^*ctrl*^, 25.3% ± 3.31% in *Tcf12*^*het*^, and 13.6% ± 2.42% in *Tcf12*^*hom*^, Figure 4G). This result parallels our ChIP-Seq analysis, which indicated a positive regulation of proliferation by TCF12. We then analyzed different stages of differentiating OLs identified by their differential expression of CC1 and OLIG1 (Marie et al., 2018; Nakatani et al., 2013). We did not detect any difference among genotypes in the proportions of recombined cells being early (CC1^+^OLIG1^-^GFP^+^ cells) and intermediate/late differentiating OLs (CC1^+^OLIG1^+^GFP^+^ cells), although we noted a trend towards more mature oligodendrocytes in *Tcf12*^*hom*^ mice (Figure 4H-I). Together, our data suggest that TCF12 is a positive regulator of OPC proliferation and likely dispensable in OL differentiation.

**Figure 4:**
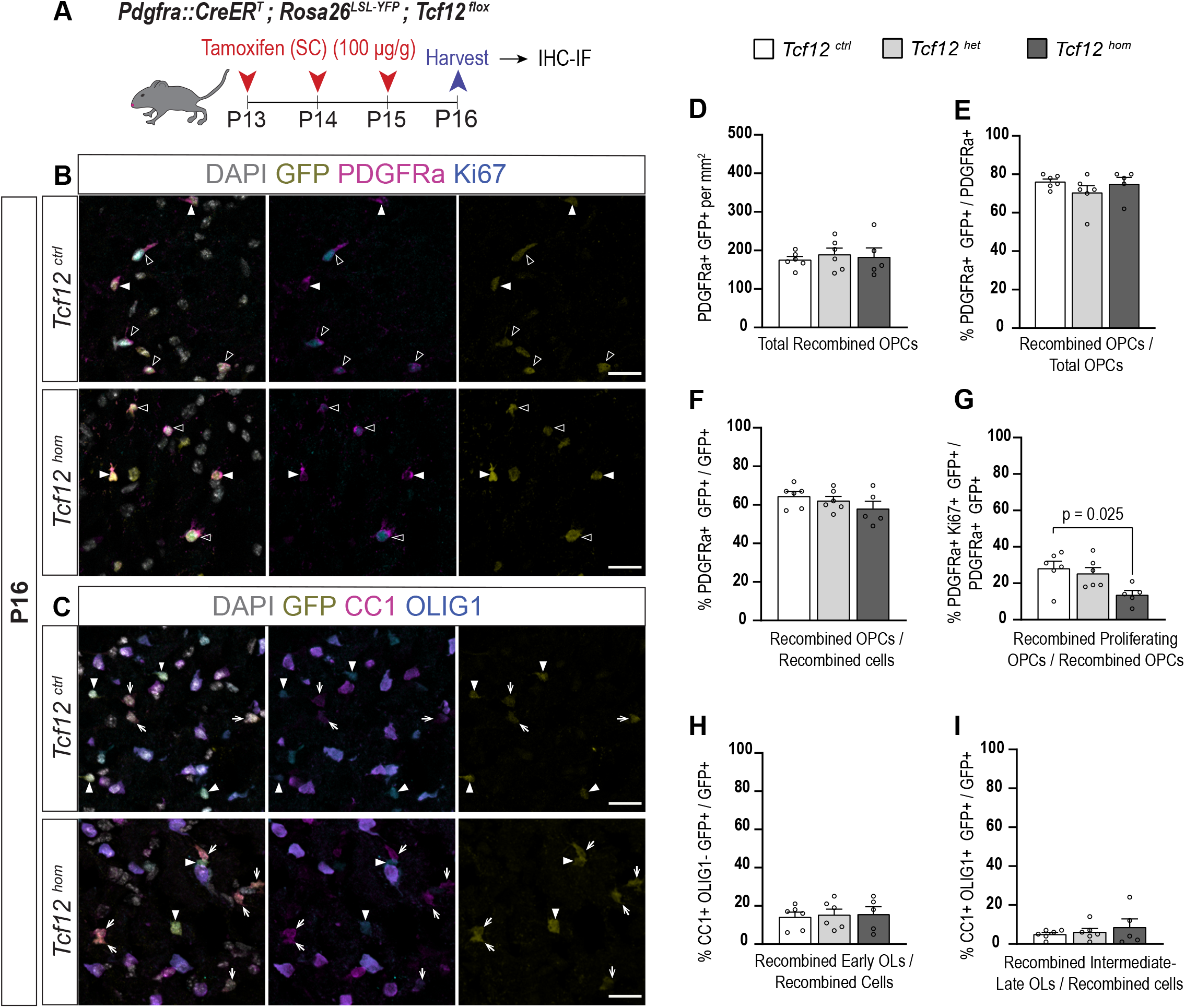
TCF12 inactivation in OPCs in vivo results in proliferation defects. **A**. Schematic representation of the experimental procedure and timeline. **B-C**. Immunostaining of YFP (recognized by an anti-GFP antibody), with PDGFRa and Ki67 (B) or CC1 and OLIG1 (C) in the corpus callosum of tamoxifen induced *Tcf12*^*ctrl*^ and *Tcf12*^*hom*^ pups at P16. Scale bars, 20μm. In B, open arrowheads show recombined proliferating OPCs (PDGFRa+ Ki67+ GFP+) and white arrowheads show non proliferating recombined OPCs (PDGFRa+ Ki67-GFP+). In C, white arrows show recombined early OLs (CC1+ OLIG1-GFP+) and white arrowheads show recombined OPCs (CC1-OLIG1+ GFP+). **D-I** Quantification of the different populations within the recombined cells (GFP+) in P16 *Tcf12*^*ctlr*^, *Tcf12*^*het*^ and *Tcf12*^*hom*^ tamoxifen induced pups. Individual points represent individual animals (n=6 *Tcf12*^*ctlr*^, n=6 *Tcf12*^*het*^ and n=5 *Tcf12*^*hom*^). Data are presented as mean + SEM. Statistical differences were evaluated with one-way ANOVA and p-values (when differences are significant) are given on the graphs.

To address whether changes in OPC proliferation and differentiation persist following *Tcf12* inactivation, we induced *Pdgfra::CreERT;Rosa*^*LSL-YFP*^*;Tcf12*^*flox*^ mice as previously at P13 and harvested them ten days later (P23; Supplementary Figure 4F-N). We noticed that the densities of recombined cells (PDGFRa+GFP+/mm2) were similar among genotypes (Supplementary Figure 4I). In addition, OPC proliferation did not change between genotypes, nor differentiating oligodendrocytes (Supplementary Figure 4J-N). These results suggest that TCF12 is, in the long term, dispensable for proper OPC proliferation and differentiation.

### Transcriptomic analyses of *Tcf12*-deficient oligodendroglial cells highlight differentiation defects and deregulation of cancer related pathways

To gain insights into the molecular mechanisms altered upon *Tcf12* inactivation, we performed a transcriptomic analysis. We purified OPCs/OLs (O4^+^ cells) at P16 from control and mutant mice three days after tamoxifen induction and performed RNA sequencing (Figure 5A). Using stringent criteria of statistical analysis (FDR <0.05, log2FC >0.5), we detected few differentially expressed genes (Supplementary Table 5). We thus analyzed the data by the method of gene set enrichment analysis (GSEA). To this goal, we first performed GSEA using a hand-curated collection of oligodendroglial gene sets derived from publications (Marques et al., 2016, 2018; Weng et al., 2019; Zhang et al., 2014) (collection provided in Supplementary Table 6). In line with our observations from immunofluorescence analysis, we observed a downregulation of gene sets related to OPCs and their proliferation in mutant animals compared to controls (Figure 5B and Supplementary Table 7). Moreover, we noted a positive enrichment of gene sets related to more differentiated oligodendrocytes in mutant compared to control animals (Figure 5B and Supplementary Table 7). Interestingly, enrichment analysis comparing *Tcf12*^*Het*^ vs *Tcf12*^*ctrl*^ cells revealed similar defects in proliferation and differentiation, although we were not able to detect these changes with the immunofluorescence approach (Supplementary Figure 5A).

**Figure 5:**
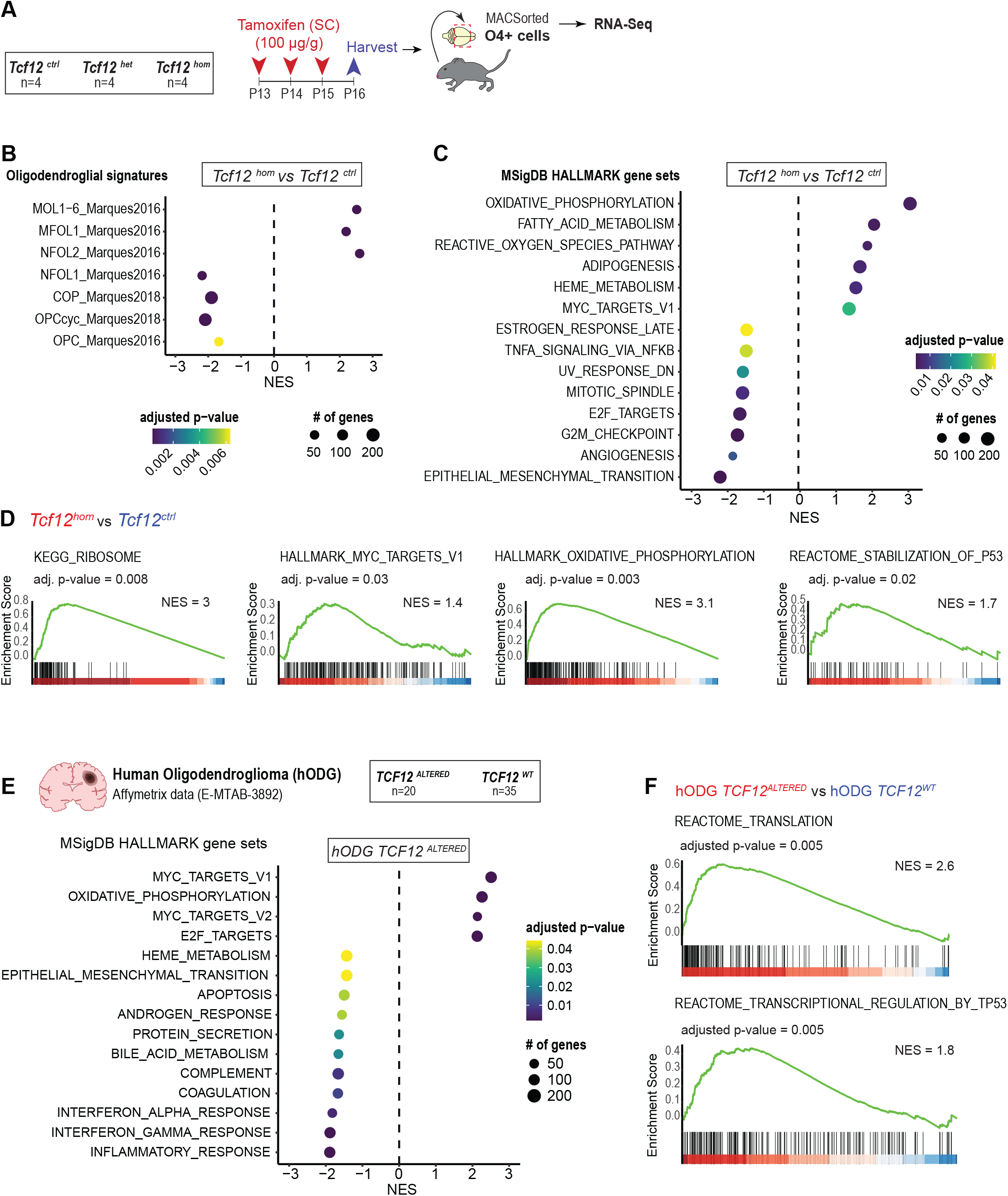
Transcriptomic analyses of *Tcf12* inactivated cells highlight defects in OPC proliferation, differentiation, and cancer-related pathways that are conserved in human gliomas. **A**. Scheme illustrating the experimental strategy for the RNA-Seq experiments. **B-C**. Gene Set Enrichment Analysis (GSEA) of *Tcf12*^*hom*^ vs *Tcf12*^*ctrl*^ O4+ cells (OPCs/OLs) using curated oligodendroglial signatures (B) and MSigDB HALLMARKS gene sets (C). **D**. GSEA plots of selected gene sets of KEGG, HALLMARKS and REACTOME collections, comparing *Tcf12*^*hom*^ vs *Tcf12*^*ctlr*^ animals. **E**. Gene Set Enrichment Analysis (GSEA) of oligodendrogliomas altered for *TCF12* (*TCF12*^ALTERED^ hODG, n=20) compared to non-altered oligodendrogliomas (*TCF12*^WT^ hODG, n=35) showing all significantly enriched gene sets of HALLMARK MSigDB collection. **F**. GSEA plots of selected gene sets for the REACTOME collection comparing *TCF12*^ALTERED^ vs *TCF12*^WT^ hODG.

To establish a broader view of the data beyond the oligodendrocyte signatures, we carried out GSEA using the highly curated HALLMARK collection from the molecular signature database (MSigDB) (Liberzon et al., 2015). In agreement with the previous analysis, comparison between *Tcf12*^*hom*^ and controls also revealed a negative enrichment of cell cycle processes (E2F targets, mitotic spindle, G2M checkpoint), together with epithelial-mesenchymal transition (EMT) gene sets, and a positive enrichment of pathways related to metabolism (oxidative phosphorylation, fatty acid, adipogenesis, and reactive oxygen species) and MYC target genes (Figure 5C). Parallel analysis of *Tcf12*^*het*^ compared to controls showed no enrichment in cell cycle processes, but upregulation of processes related to cholesterol homeostasis, and downregulation of immune signatures (inflammatory and interferon responses), EMT and NOTCH signaling (Supplementary Figure 5B). Querying additional collections from the molecular signature database (REACTOME, KEGG, GO_Biological_Process), we observed that *Tcf12* inactivation was also associated with negative regulation of developmental pathways (such as NOTCH, BMP) and pathways related to cancer (such as cell cycle, extracellular matrix, ribosome, TP53,) (Supplementary Figure 5C). Interestingly, the absence of functional TCF12 induced a deregulation of ribosome biogenesis and translation-related pathways (Figure 5D, Supplementary Figure 5C) which are a strictly tuned in a multi-staged process controlling diverse cellular responses, such as cell proliferation and growth (Hetman & Slomnicki, 2019).

### Conservation of TCF12 regulated pathways in human gliomas

To determine how TCF12-dependent pathways may be relevant in gliomas, we took advantage of our previously published cohort of *TCF12*-mutated oligodendrogliomas (Labreche et al., 2015). We re-analyzed the transcriptomics data, comparing samples with *TCF12* alterations (mutations and/or loss of heterozygosity and/or copy number loss of heterozygosity without loss of heterozygosity; n=20) to non-altered (non-mutated and normal TCF12 genomic status; n=35) samples (Supplementary Table 8). We performed gene set enrichment analysis using the HALLMARK collection from the molecular signature database. Interestingly, this analysis showed positive enrichment of pathways related to MYC, oxidative phosphorylation, cell cycle (E2F targets) in *TCF12*-altered tumors, similar to our analysis of *Tcf12*-inactivated cells (Figure 5E). In addition, EMT and immune response signatures were negatively enriched in *TCF12*-altered tumors. We also interrogated the REACTOME collection and found that pathways related to translation and regulation of p53 activity were positively enriched in *TCF12-*altered tumors (Figure 5F). These data indicate that the TCF12-regulated processes that we identified from our molecular analyses of mouse oligodendroglial cells are conserved in a glioma setting.

## Discussion

Oligodendrocyte precursor cells (OPCs) have been proposed to be the cells of origin for gliomas and responsible for tumor expansion (C. Liu et al., 2011; Persson et al., 2010; Sugiarto et al., 2011; Weng et al., 2019), although mutations may arise as early as the neural stem cell stage (J. H. Lee et al., 2018; C. Liu et al., 2011). Understanding how genes altered in gliomas impact the lineage of origin can provide insights into their implication in gliomas, as these functions may be conserved in a tumor context. In this study, we aimed to explore the roles of TCF12 in the oligodendroglial lineage and in gliomagenesis. Although TCF12, similar to other members of the E protein family (TCF3, TCF4), is a ubiquitously expressed protein, its function remains poorly characterized in cell-specific contexts. *Tcf12* was reported to be expressed in spinal cord oligodendroglial cells and oligodendroglial cultures (Fu et al., 2009; Sussman et al., 2002; Wedel et al., 2020). Here, querying resources of bulk and single cell transcriptomic data sets from embryonic and postnatal mouse brain (Marques et al., 2016, 2018; Zhang et al., 2014), we find that *Tcf12* is expressed throughout oligodendrocyte lineage development, with higher transcript levels in progenitor and cells at early stages of differentiation. By immunofluorescence, we demonstrate a similar pattern of TCF12 protein in postnatal and adult oligodendroglia, with TCF12 levels peaking in OPC and differentiating oligodendrocytes.

We report the first analysis of TCF12 genome wide binding sites in oligodendroglial cells acutely isolated from the postnatal cerebral cortex. TCF12 binding sites were enriched in promoter regions compared to other genomic regions, and associated with active histone marks, suggesting that TCF12 acts as a transcriptional activator in oligodendrocyte lineage cells. The key regulators of oligodendrocyte development ASCL1 and OLIG2 are known to dimerize with E-proteins to control gene expression. TCF12 binding sites partially overlap with those of ASCL1 and OLIG2 (C Marie and C Parras, unpublished data) suggesting interaction between TCF12 and these proteins in transcriptional regulation of oligodendroglial cells. Interestingly, a recent study pointed to a preferential interaction of TCF12 with OLIG1 in HEK cells (Wedel et al., 2020). Future studies will be needed to determine the contribution of the different TCF12 heterodimers in the regulation of OPC proliferation and differentiation.

We further show that specific inactivation of *Tcf12* in postnatal OPCs result in a significant decrease in the proportion of proliferating OPCs without impacting the proportions of OPCs and oligodendrocytes. This result indicates that TCF12 controls OPC proliferation, as suggested by our ChIP-Seq data and in agreement with our observations of high *Tcf12* expression in neural stem and progenitors of the mouse brain. Transcriptomic analyses of *Tcf12* deficient cells offered a higher resolution of altered processes and further pointed out differentiation defects, that we were unable to detect from our immunohistochemical analysis, although we noted a trend towards more mature oligodendrocytes in *Tcf12*-deficient mice. Enrichment of oligodendrocyte signatures in *Tcf12*-deficient cells may reflect the consequences of decreased OPC proliferation. Our data thus suggest that in the oligodendroglial lineage, TCF12 may primarily act on proliferation, rather than differentiation. This is in agreement with a recent study showing that ectopic TCF12 does not affect oligodendrocyte differentiation of brain organotypic slices (Wedel et al., 2020). However, our finding of TCF12 binding to the promoters of oligodendrocyte genes implies a possible and subtle implication of TCF12 in the control of differentiation. We noted that the effects of *Tcf12* inactivation on proliferation are transient. Given that all three E-protein genes (*Tcf12, Tcf3, Tcf4*) are expressed in developing oligodendrocytes, it is possible that TCF3 and TCF4 may compensate for TCF12 deficiency, as shown for *Tcf12-/-* mouse cerebella that display upregulation of *Tcf4* transcripts (Ravanpay & Olson, 2008). Although we did not detect an upregulation in *Tcf3* and *Tcf4* transcripts in *Tcf12*-deficient OPCs three days post inactivation (Supplementary Table 5), we cannot exclude that a compensation takes place later on, given the strong levels of *Tcf4* transcripts present in OPCs.

Transcriptomic analysis of both *Tcf12*-deficient oligodendroglia (this study) and TCF12-mutated gliomas (Labreche et al., 2015) revealed few significantly differentially expressed genes in the context of TCF12 inactivation compared to wild type TCF12. This result is intriguing given that we find TCF12 binds over 15,000 genes in the genome of oligodendroglial cells and therefore one might expect substantial deregulation of gene expression in the absence of functional TCF12. However, the apparent lack of gene expression differences may in fact highlight the diversity of processes directly controlled by TCF12, as many of them may interact and regulate each other, leading to an overall normalization of gene expression.

Interestingly, our ChIP-Seq and RNA-Seq data suggest a control by TCF12 of several processes involved in cancer, many of which are also perturbed in TCF12 altered oligodendrogliomas. For example, we observed a negative enrichment of terms related to the extracellular matrix and epithelial-mesenchymal transition (EMT) in *Tcf12-*deficient OPCs and in *TCF12*-altered oligodendrogliomas. Accordingly, TCF12 was shown to repress E-cadherin expression, and its expression has been correlated with increased invasion, migration, and metastasis in several cancers (He et al., 2016; C.-C. Lee et al., 2011; Luo et al., 2020), including GBM (Zhu et al., 2021). An important difference we noted between our mouse model and *TCF12*-altered oligodendrogliomas is that while proliferation is decreased in *Tcf12*-deficient OPCs, loss of *TCF12* is associated with increased proliferation in human oligodendrogliomas. We previously showed that *TCF12* mutations in oligodendrogliomas were associated with more aggressive tumor features (Labreche et al., 2015), indicative of a tumor-suppressive function for TCF12. In line with these findings, TCF12 was recently identified as a master regulator of the differentiated, rather than stem-like, state in glioblastomas (Castellan et al., 2021). In contrast, a previous study showed that *TCF12* silencing decreased proliferation and invasion in glioma cell lines (Godoy et al., 2016). All these data suggest that mechanisms induced by the mutational and cellular contexts in gliomas may interfere with TCF12-mediated regulation of proliferation.

Importantly, our study implies novel roles for TCF12 in the control of ribosome biogenesis and translation. TCF12 has been shown to bind the bHLH transcription factor MYC in rat fibroblasts (Agrawal et al., 2010). MYC directly controls ribosome biogenesis and translation, by inducing the transcription of ribosomal RNA, ribosomal proteins and genes involved in the maturation of ribosomal RNAs (Piazzi et al., 2019). Perturbation of the ribosome biogenesis process has been shown to activate p53 (Piazzi et al., 2019). Interestingly, processes related to MYC, TP53 and ribosome/translation were enriched in both *Tcf12-*deficient oligodendroglial cells and *TCF12*-altered oligodendrogliomas. In addition, we detected binding of TCF12 on promoters of *Myc, Mycn* and genes involved in ribosome biogenesis (such as *Rpl5, Ruvbl2, Fbl*) in oligodendroglial cells. Our study is the first to suggest a link between TCF12 and ribosome biogenesis. Of note, interactions between E-proteins and ribosome biogenesis were previously reported: TCF4 is detected in the nucleolus, where ribosome biogenesis occurs, and loss of function mutants inhibit protein synthesis in rat hippocampal neurons (Slomnicki et al., 2016). Moreover, TCF4 overexpression represses MYC target genes in a leukemic cell line (N. Liu et al., 2019). In conclusion, our study suggests that TCF12 directly regulates many signaling pathways in oligodendroglia development that are relevant in a glioma context.

## Author contributions

SA and EH conceptualized the study. SA, CM, CP and EH designed experiments and analyzed data. SA and CM performed experiments. BG, JG and CP analyzed transcriptomic data generated in this paper and from public databases. MS, CP and EH obtained funding. EH supervised the study. SA, CP and EH wrote the manuscript. All authors revised the manuscript.

## Acknowledgements

We acknowledge funding from Ligue Nationale contre le Cancer (to MS), Fondation ARC (PJA 20151203259 to EH), National Multiple Sclerosis Society (NMSS RG-1501-02851 to CP), and the Fondation pour l’Aide à la Recherche sur la Sclérose en Plaques (ARSEP 2015, 2018, 2019 to CP), Fondation pour la Recherche Médicale (to CP). SA is recipient of scholarships from Ligue Nationale Contre le Cancer and Fondation pour la Recherche Médicale. We acknowledge the contribution of SiRIC CURAMUS (INCA-DGOS-Inserm_12560) which is financially supported by the French National Cancer Institute, the French Ministry of Solidarity and Health and Inserm. The research leading to these results has received funding from the program “Investissements d’avenir” ANR-10-IAIHU-06. Part of this work was carried out on the iGenSeq, CELIS, Histomics, PHENOPARC, Icm.Quant and Data and Analysis core facilities of ICM. We gratefully acknowledge Yannick Marie for assistance on RNA sequencing. We thank Isabelle Le Roux for critical input and reading of the manuscript.

## Data Availability Statement

ChIPseq and RNAseq data generated in this study are available through the Gene Expression Omnibus (*data will be deposited upon acceptance of the manuscript*). The data that support the findings of this study are available from the corresponding author upon reasonable request.

## Figure Legends

Supplementary Figure 1 (related to Figure 1)

**A**. OncoPrint showing *TCF12* alterations along with other genes commonly altered in gliomas (TCGA GBM/LGG cohort). Each column represents one sample. Only the *TCF12* altered samples (210/1106) are displayed. The cumulative frequency of alterations of each gene observed in the total number of patient samples is indicated.

**B**. Chart summarizing the estimated co-occurrence of *TCF12* alterations with genes commonly altered in gliomas (from Supplementary Figure 1A). Statistical significance was determined with Fisher’s exact test. Analysis was performed using the “Mutual Exclusivity” tool by “cbioportal”.

Supplementary Figure 2 (related to Figure 2)

**A**. UMAP representing the compilation of the two single cell RNA-Seq data sets by Marques *et al* 2016 & 2018. **B**. UMAP with colored dots representing the 9 stages of differentiation (simplified clusters), from neural stem cells to mature myelinating oligodendrocytes. **C**. Dot plot summarizing the average mRNA expression of the three E proteins (*Tcf12, Tcf3* and *Tcf4*) and the percentage of expressing cells in each one of the clusters of the Supplementary Figure 2B.

Supplementary Figure 3 (related to Figure 3)

**A**. Scheme illustrating the experimental strategy for the ChIP-Seq experiments. **B**. Quantification of PDGFRa+ and CNP+ cells (as percentage of the total cells) of the immunofluorescence immunocytochemistry of cells that were plated directly after the MACSorting and kept in culture for 2 hours. PDGFRa+ cells are OPCs. CNP+ cells are mostly OLs (with some OPCs expressing CNP). Data are presented as mean + SEM. Individual points correspond to individual animals (n=6 wild type mice). **C**. Pie charts showing the distribution of TCF12-bound sites between promoter (left) and enhancer (right) regions in NPCs and in association with epigenetic marks (H3K4me3 and H3K27Ac = active, H3K4me = poised, H3K27me3= repressed; NA= no epigenetic mark). **E**. Selected ChIP-Seq tracks for TCF12, input control and epigenetic marks in promoter regions of *Mycn, Myc, Rpl5, Ruvbl2, Fbl* in OPCs/OLs.

Supplementary Figure 4 (related to Figure 4)

**A**. Illustration of the genetics of the mouse lines used to create the *Pdgfra-CreER*^*T*^; *Rosa26*^*LSL-YFP*^; *Tcf12*^*flox*^ mice. **B**. Genetics of the animals used for the experiments. **C**. Experimental procedure for the validation of the mouse model. **D**. RT-qPCR analysis of *Tcf12* expression in O4+ MACSorted cells from P16 *Tcf12*^*ctlr*^, *Tcf12*^*het*^ and *Tcf12*^*hom*^ induced pups. Individual points correspond to individual animals (n=3 *Tcf12*^*ctlr*^, n=3 *Tcf12*^*het*^ and n=4 *Tcf12*^*hom*^). Data are presented as mean + SEM. Individual points represent individual animals. Statistical differences were evaluated with one-way ANOVA and p-values are given on the graphs. **E**. Quantification of GFP+, PDGFRa+ and OLIG2+ cells plated directly after the MACSorting and kept in culture for 2 hours. GFP+ cells are considered as recombined cells. PDGFRa+ cells are OPCs. OLIG2+ cells correspond to OPCs and OLs. Data are presented as mean + SEM. Individual points correspond to individual animals (n=4 *Tcf12*^*ctlr*^, n=3 *Tcf12*^*het*^ and n=3 *Tcf12*^*hom*^). Differences were analyzed with two-way (% total cells) or one-way (% recombined cells) ANOVA. No statistically significant differences were noted. **F**. Schematic representation of the experimental procedure and timeline for the P23 timepoint. **G-H**. Immunostaining of YFP (recognized by an anti-GFP antibody), with PDGFRa and Ki67 (B) or CC1 and OLIG1 (C) in the corpus callosum of tamoxifen induced *Tcf12*^*ctlr*^ and *Tcf12*^*hom*^ pups (P23). Scale bars, 20μm. B: open arrowheads show recombined proliferating OPCs (PDGFRa+ Ki67+ GFP+) and white arrowheads show non proliferating recombined OPCs (PDGFRa+ Ki67-GFP+). C: white arrows show recombined OLs (CC1+ OLIG1+ GFP+) and white arrowheads show recombined OPCs (CC1-OLIG1+ GFP+). **I-N**. Quantification of the different populations within the recombined cells (GFP+) in P23 *Tcf12*^*ctlr*^, *Tcf12*^*het*^ and *Tcf12*^*hom*^ tamoxifen induced pups. Data are presented as mean + SEM. Individual points represent individual animals (animals quantified I-K: n=5 *Tcf12*^*ctlr*^, n=5 *Tcf12*^*het*^ and n=4 *Tcf12*^*hom*^, animals quantified L-N: n=4 *Tcf12*^*ctlr*^, n=4 *Tcf12*^*het*^ and n=4 *Tcf12*^*hom*^). Statistical differences were evaluated with one-way ANOVA. No statistically significant differences were noted.

Supplementary Figure 5 (related to Figure 5)

**A-B**. Gene Set Enrichment Analysis (GSEA) of *Tcf12*^*het*^ vs *Tcf12*^*ctrl*^ O4+ cells (OPCs/OLs) using curated oligodendroglial signatures (A) and MSigDB HALLMARKS gene sets (B). **C**. Bar plots of GSEA normalized enrichment score (NES) of selected gene sets (positively and negatively enriched) from the GO_Biological_Process, KEGG and REACTOME collections of the MSigDB, comparing *Tcf12*^*hom*^ vs *Tcf12*^*ctrl*^ animals. Adjusted p-values are given on the corresponding bars.

